# Oxytocin attenuates microglial activation and restores social and non-social memory in the APP/PS1 mouse model of Alzheimer’s disease

**DOI:** 10.1101/2022.05.07.491031

**Authors:** Maria Clara Selles, Juliana T.S. Fortuna, Yasmin P.R. de Faria, Beatriz Monteiro Longo, Robert C. Froemke, Moses V. Chao, Sergio T. Ferreira

## Abstract

Alzheimer’s disease (AD) is the main cause of dementia in the elderly and is characterized by memory loss, social withdrawal and neurodegeneration, eventually leading to death. Brain inflammation has emerged as a key pathogenic mechanism in AD. We hypothesized that oxytocin, a pro-social hypothalamic neuropeptide with anti-inflammatory properties, could have therapeutic actions in AD. We investigated oxytocin production in mouse models of AD, and evaluated the therapeutic potential of intranasal oxytocin. We observed lower levels of hypothalamic oxytocin in wild-type mice following brain infusion of amyloid-β oligomers (AβOs), as well as in APP/PS1 AD model mice. Treatment of APP/PS1 mice with intranasal oxytocin reduced microglial activation and favored deposition of Aβ in dense core plaques, a potentially neuroprotective mechanism. Oxytocin further alleviated social and non-social memory impairments in APP/PS1 mice. Our findings point to oxytocin as a potential therapeutic target to reduce brain inflammation and correct memory deficits in AD.

## Introduction

Dementia affects approximately 50 million people worldwide with nearly 10 million new cases per year, and AD is responsible for 60-70% of those cases (*1*). Among other features, AD is characterized by brain accumulation of the Aβ peptide and by prominent brain inflammation, comprising aberrant production and release of pro-inflammatory cytokines, and activation of astrocytes and microglia to toxic phenotypes (*2,3*). The inflammatory hypothesis for AD (*4*) has stimulated a large number of recent studies investigating the role of brain inflammation in AD pathogenesis and cognitive dysfunction (*5,6*). In particular, studies have aimed to attenuate microglial activation in an attempt to arrest or delay the development of the disease (*5,7,8*).However, despite encouraging results in preclinical studies (*2*), none of those potential treatments has reached or been successful in clinical trials to date (*9*). Thus, it is critical to identify therapeutic agents that can regulate microglial activation and inflammation, to prevent or reduce disease progression in Alzheimer’s patients.

A promising anti-inflammatory therapeutic molecule is oxytocin, a nonapeptide hormone produced in the supraoptic nucleus and paraventricular nucleus (PVN) of the hypothalamus (*10*). In the periphery, oxytocin is well-known for its role in lactation and parturition (*10*). In the brain, oxytocin plays important roles in social behavior, including bonding and reproduction (*11*), and recent studies have described its neuroprotective and anti-inflammatory actions (*12,13*). Moreover, Ye et al. pointed oxytocin as an intervention for early AD that can modulate the immune response in AD mice and prevent cognitive decline (*14*). As a role for oxytocin in AD-related neuroinflammation has not been examined once the pathology is already stablished, we hypothesized that oxytocin treatment could attenuate microglial activation and reverse memory deficits in aged AD mouse models.

## Results

### Oxytocin production is down-regulated in AD models

We first examined whether oxytocin production was affected in AD models. Oxytocin expression was reduced in mouse hypothalamic slices exposed to AβOs (Fig. 1A), toxins that accumulate in AD brains and are proximally implicated in pathogenesis (*15,16*). Reduced hypothalamic expression of oxytocin was further verified *in vivo* in 3 month old wild-type male mice that received intracerebroventricular infusions of AβOs for 5 consecutive days, as well as in aged male APPswe/PS1ΔE9 (APP/PS1) mice (a genetic model of AD) (Fig. 1B and C). These findings are in agreement with a report of downregulated oxytocin mRNA in young (4 month-old) APP/PS1 mice (*17*), and suggest that oxytocin production is defective in AD models.

**Fig. 1.**
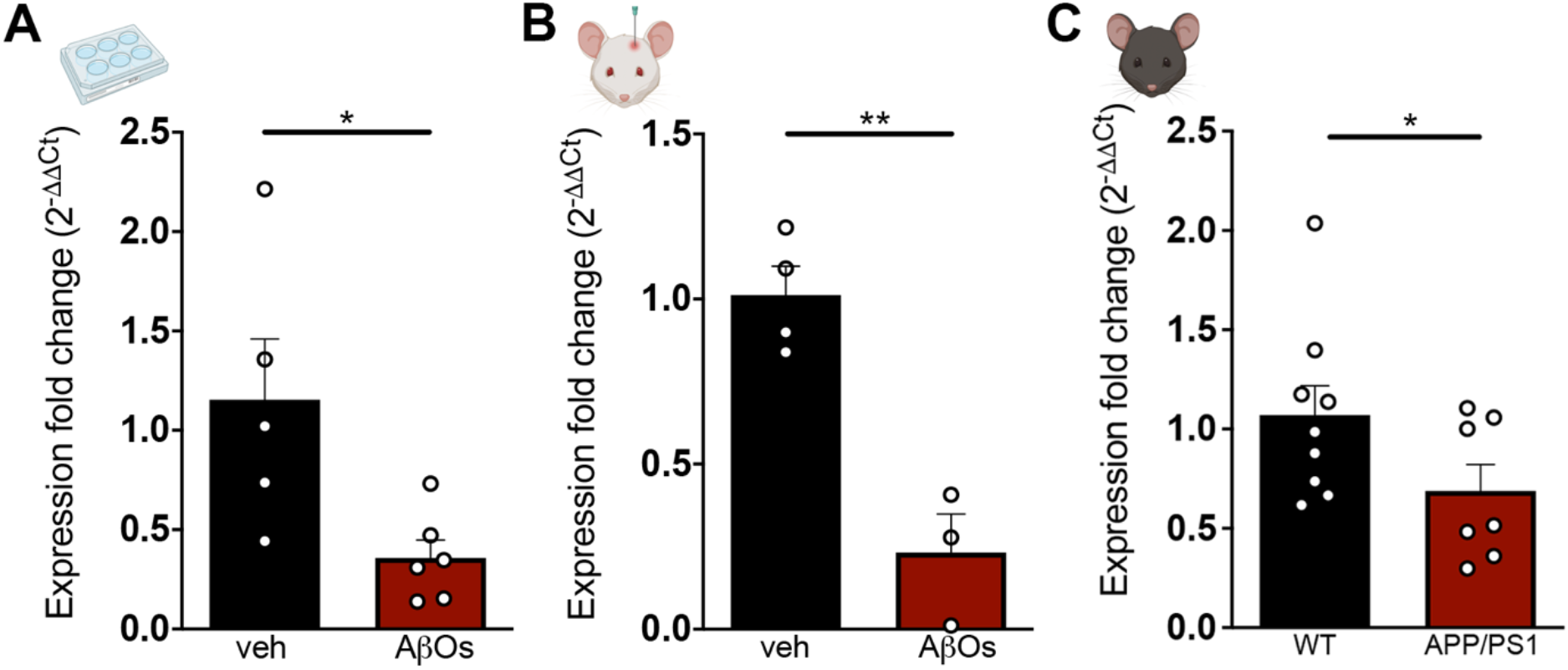
Hypothalamic expression of oxytocin is down-regulated in AD models. Oxytocin expression was determined by qPCR in (**A**) acute hypothalamic slices (from 3 month old wild-type male mice) exposed to AβOs (0.5 μM) or vehicle for 3h; (**B**) 3 month old wild-type male mice that received intracerebroventricular infusions of AβOs (10 pmol) or vehicle for 5 consecutive days; and (**C**) 12-13.5 month old male wild-type and APP/PS1 mice. Symbols represent data from individual mice, unpaired one-tailed Student’s t-test, *p <0.05, **p<0.01.

### Chronic intranasal oxytocin increases hippocampal oxytocin levels

To test the hypothesis that reduced brain oxytocin could be a contributing factor to pathogenesis in AD mice, we developed a protocol to treat mice with intranasal oxytocin aiming to increase oxytocin levels in the brain. We treated 9-10 month old C57Bl/6 wild-type mice intranasally with 8 ng oxytocin three times a week for 6 weeks (Fig. 2A), and observed a ∼70% increase in hippocampal oxytocin peptide levels (Fig. 2B). In addition, and consistent with decreased fear response induced by oxytocin (*18*), mice treated intranasally with oxytocin showed decreased freezing behavior in a fear conditioning paradigm (Fig. 2C). These results show that intranasal administration of oxytocin effectively increases oxytocin levels in the mouse hippocampus.

**Fig. 2.**
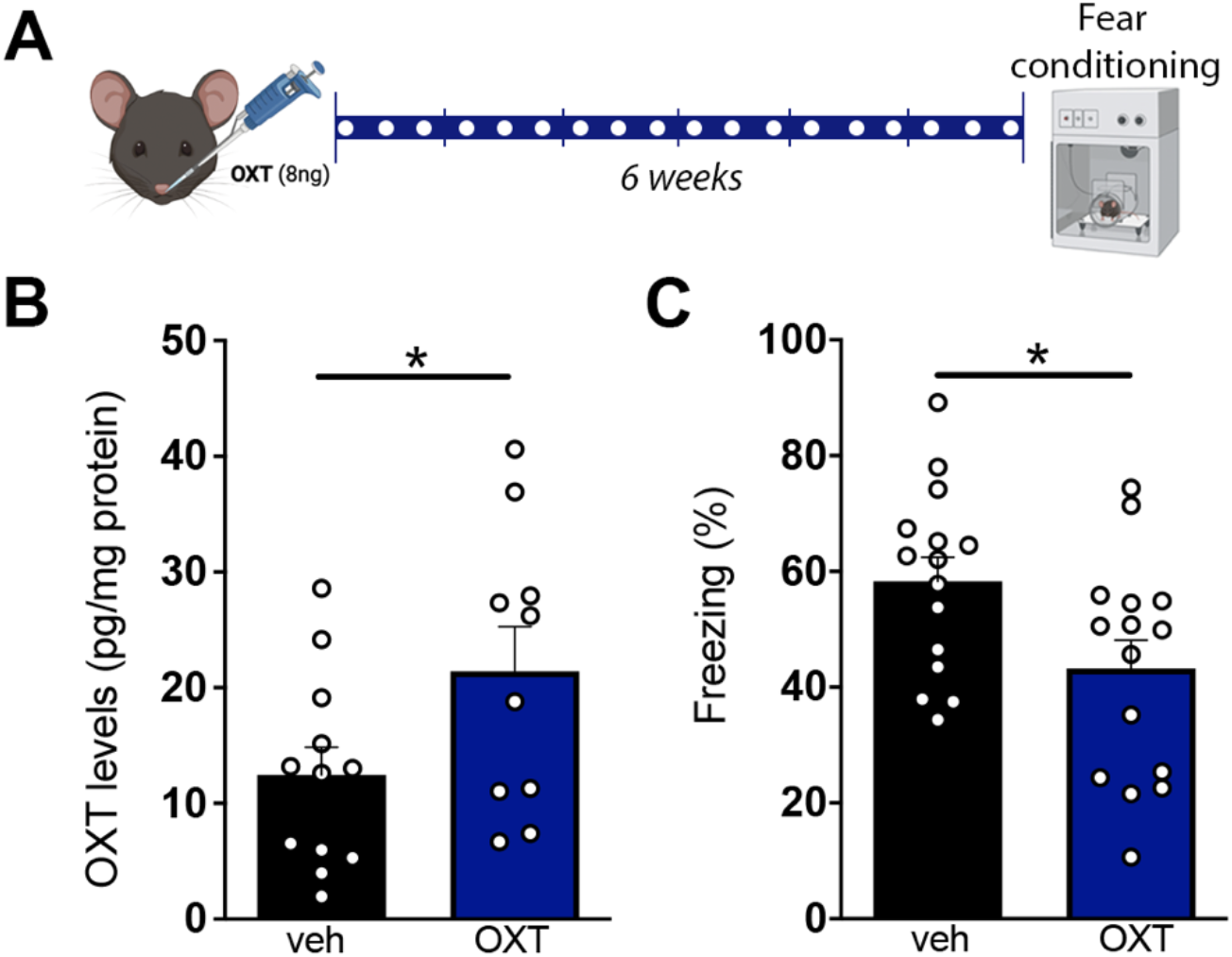
Chronic intranasal administration of oxytocin increases hippocampal oxytocin. (**A**) Experimental design used for intranasal treatment with oxytocin. White dots indicate intranasal administration of vehicle (veh) or oxytocin (OXT). (**B**) Hippocampal oxytocin levels (OXT) in 9-10 months old wild-type male mice treated intranasally with veh or OXT. (**C**) Effect of intranasal treatment with veh or OXT on freezing behavior in the contextual fear conditioning paradigm. Symbols represent individual mice. Unpaired one-tailed Student’s t-test.

### Intranasal oxytocin restores hypothalamic oxytocin in APP/PS1 mice

We next determined oxytocin levels within hypothalamic PVN oxytocinergic neurons of wild-type and APP/PS1 mice treated intranasally with oxytocin (Fig. 3A). In harmony with reduced mRNA levels reported above (Fig. 1C), we observed decreased hypothalamic oxytocin peptide immunoreactivity in APP/PS1 mice compared to wild-type littermates. Surprisingly, oxytocinergic cells from APP/PS1 mice treated with exogenous oxytocin exhibited higher oxytocin immunoreactivity (Fig. 3B). These results raise the possibility that the increase in hippocampal oxytocin after intranasal treatment with oxytocin could, at least in part, be due to stimulation of endogenous hypothalamic production of oxytocin.

**Fig. 3.**
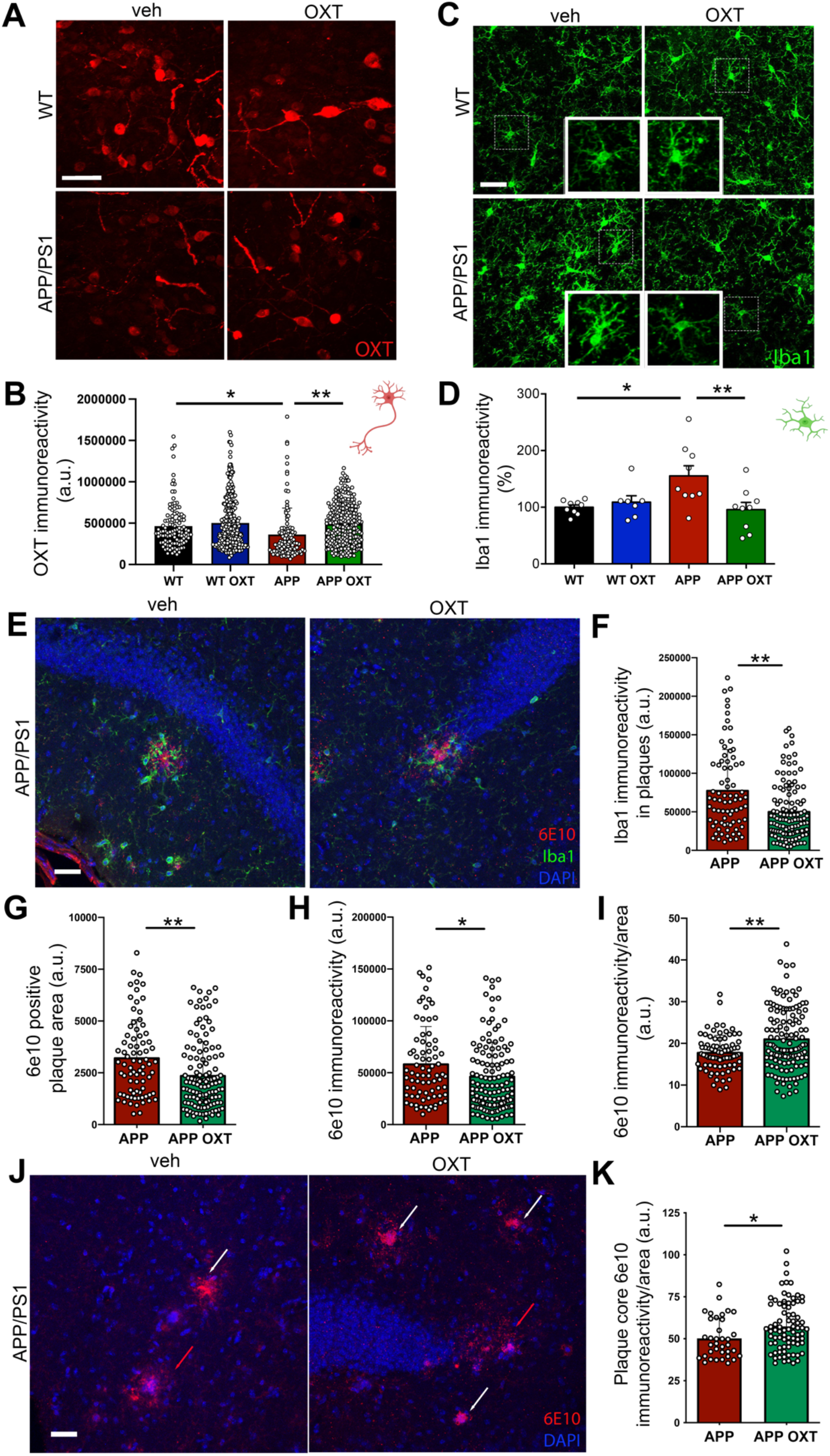
Cellular and molecular impact of intranasal oxytocin in APP/PS1 mouse brains. (**A**) Representative images for oxytocin (OXT) immunohistochemistry in the PVN of wild-type (WT) or APP/PS1 (APP) mice treated with intranasal vehicle (veh) or OXT. (**B**) Quantification of OXT immunoreactivities. Symbols represent individual cells. N = 2-3 mice per group; two-way ANOVA followed by Dunnett’s post-hoc test. (**C-F**) Effects of oxytocin on microglia. (**C**) Representative images for Iba1 immunohistochemistry in the CA1 region of the hippocampus of wild-type or APP/PS1 mice treated with intranasal OXT or vehicle. (**D**) Quantification of Iba1 immunoreactivity. Symbols represent mean values from individual mice; two-way ANOVA followed by Dunnett’s post-hoc test. (**E**) Representative double immunofluorescence image showing amyloid plaques (6E10 immunoreactivity, red) and Iba1 immunoreactivity (green) in the hippocampus of an APP/PS1 mouse. (**F**) Iba1 immunoreactivity surrounding plaques. (**E**,**G**-**I**) Effects of OXT on amyloid plaques. (**G**) 6E10-positive area of individual plaques, (**H**) 6E10 immunofluorescence intensity per plaque, (**I**) 6E10 immunofluorescence intensity normalized per plaque area. (**J**,**K**) Effect of oxytocin on plaque cores. Plaque core 6E10 immunofluorescence/area ratio was quantified. (**J**) Immunofluorescence (6E10, red) showing diffuse (6E10 immunofluorescence/area ratio < 35 a.u., red arrows) and dense core (6E10 immunofluorescence/area ratio > 35 a.u., white arrows) plaques in the hippocampi of APP/PS1 mice that received intranasal veh or OXT. (**K**) Quantification of plaque core density (6E10 immunofluorescence intensity normalized per plaque core area) in dense core plaques. Symbols represent individual plaques. N = 3-5 mice per group; unpaired two-tailed Student’s t-test, *p<0.05, **p<0.01. Scale bars: 50 μm.

### Intranasal oxytocin attenuates microglial activation in the hippocampus of APP/PS1 mice

While previous studies have investigated the regulation of neuronal function by oxytocin signaling (*11,19*), much less is known on the effects of oxytocin on other cell types of the central nervous system, including microglia. Yuan et al. (2016) showed that oxytocin prevents LPS-mediated microglial activation *in vitro* and *in vivo* (*12*). That study was followed by description of the anti-inflammatory actions of oxytocin in neurological conditions, including autism, post-traumatic stress disorder and early AD (*14, 20-22*). Because microglial activation is thought to play a major role in AD pathogenesis, we tested whether oxytocin might improve behavioral deficits in APP/PS1 mice by attenuating microglia-mediated brain inflammatory responses.

To test this hypothesis, we assessed the effect of intranasal treatment with oxytocin on the activation state of microglia in mice. In line with a recent report (*23*), we first confirmed that microglia exhibited increased Iba1 immunoreactivity (indicative of activation) in the hippocampi of aged APP/PS1 mice compared to wild-type controls, and found that treatment with intranasal oxytocin reduced microglial activation in APP/PS1 mice (Fig. 3C and D). Further, Iba1 immunoreactivity surrounding hippocampal amyloid plaques was decreased in APP/PS1 mice treated with oxytocin compared to vehicle treatment (Fig. 3E and F).

### Intranasal oxytocin alters amyloid plaque morphology in the hippocampus of APP/PS1 mice

To evaluate consequences of oxytocin treatment on Aβ accumulation, we quantified amyloid plaque area and immunoreactivity using the Aβ-specific 6E10 antibody in the hippocampi of APP/PS1 mice. We observed smaller amyloid plaques and decreased total Aβ immunoreactivity in the hippocampi of mice treated with oxytocin compared to vehicle-treated controls (Fig. 3G and H). Interestingly, however, plaques in oxytocin-treated mice had higher densities of Aβ immunoreactivity per unit area (Fig. 3I). Huang and collaborators (2021) recently reported results suggesting that microglia act as “trash compactors” and promote dense core plaque formation (*24*), a mechanism that might decrease soluble Aβ levels and be neuroprotective in AD (*25*). This prompted us to evaluate Aβ immunoreactivities within plaque cores. For this analysis, we calculated core density as the ratio between 6E10 immunoreactivity and area in plaques presenting a dense core (core 6E10 immunoreactivity/area ratio greater than 35 arbitrary units (a.u.)). We found that APP/PS1 mice treated with oxytocin had higher density plaque cores compared to vehicle-treated mice (Fig. 3J and K). These results suggest that oxytocin treatment favors Aβ deposition in the form of dense core plaques.

### Oxytocin reduces AβO-induced microglial activation in vitro

To determine whether the attenuation of microglial activation by oxytocin observed *in vivo* reflected a direct regulatory action of oxytocin on these cells, we isolated and cultured microglial cells *in vitro* (*26*). We observed that treatment with oxytocin (10 nM) prevented microglial activation induced by AβOs (500 nM) (Fig. 4A and B). AβOs induced an increase in Iba1 immunoreactivity and a decrease in the number of microglial processes, consistent with induction of a phagocytic microglial phenotype. Treatment with oxytocin blocked the increase in Iba1 immunoreactivity, and attenuated the decrease in microglial ramification in five out of six independent cultures evaluated (Fig. 4A and C). These results indicate that oxytocin attenuates microglial activation by AβOs.

**Fig. 4.**
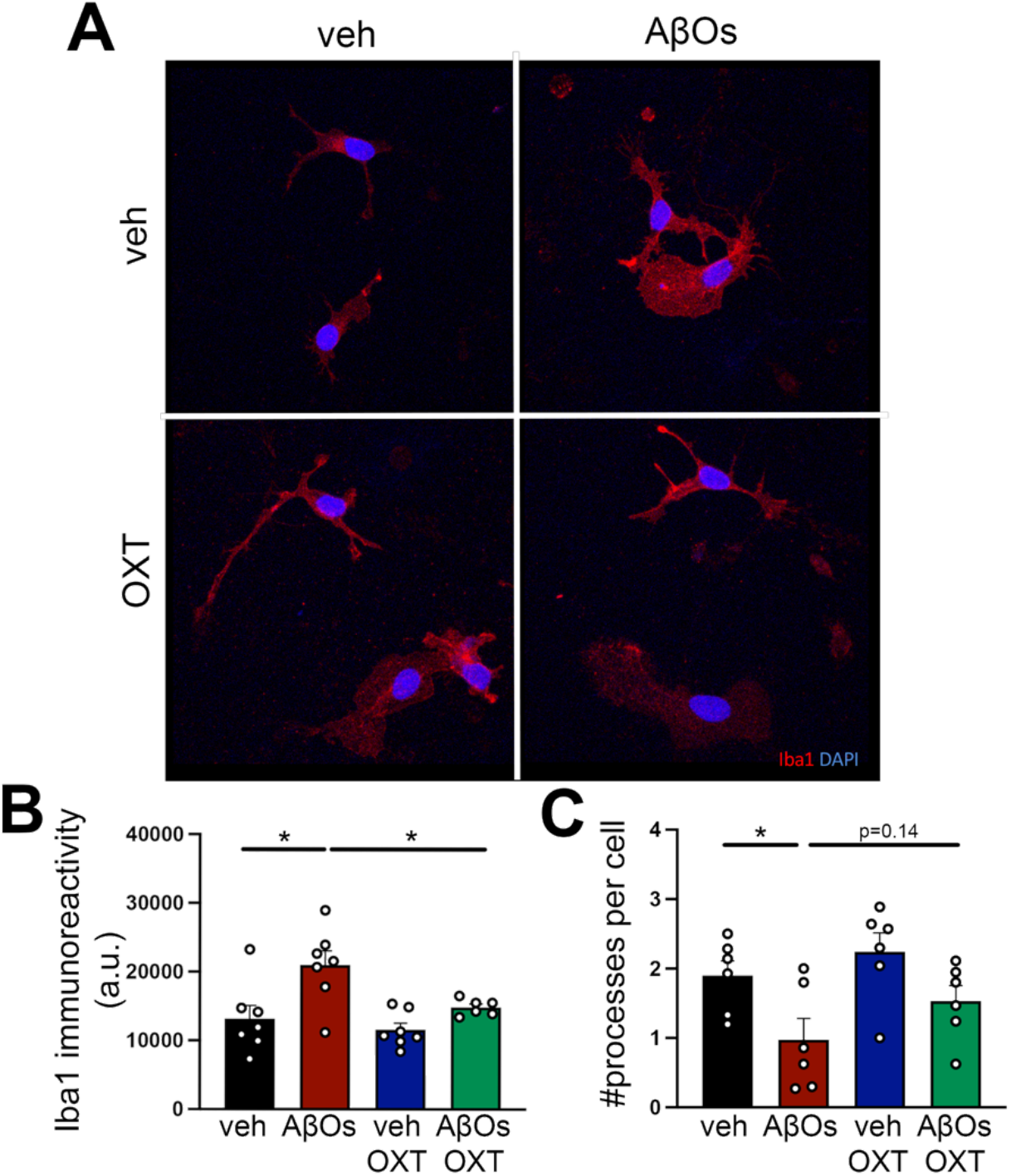
Oxytocin attenuates AβO-induced microglial activation *in vitro*. Microglia were immunopanned from P18 rat pups, kept in culture for 5 DIV and treated with oxytocin (OXT, 10 nM) and/or AβOs (500 nM) for 3h. (**A**) Representative Iba1 immunocytochemistry images. (B), Iba1 immunoreactivity and (C) number of primary microglial processes per cell. N = 6-7 independent cultures. Mixed model (**B**) and (**C**) Repeated measures ANOVA followed by Dunnett’s post-hoc test. Symbols represent mean values from independent microglial cultures. *p<0.05.

### Intranasal treatment with oxytocin reverses social, object recognition and spatial memory impairments in APP/PS1 mice

Finally, we tested whether oxytocin could comprise a therapeutic strategy for AD by assessing the effects of intranasal oxytocin on memory in APP/PS1 mice. We first studied the effect of intranasal treatment with oxytocin on social memory in APP/PS1 mice. For this, we used a 5-trial social memory test, which evaluates habituation to an intruder mouse during the first 4 trials, and novelty response to a novel intruder on the 5th trial (*27*) (Fig. 5A). While wild-type animals showed the expected decrease in exploration times across trials 1-4, APP/PS1 mice did not (Fig. 5B,C). In addition, APP/PS1 mice did not exhibit increased exploration in trial 5, when a novel mouse was presented to the test mouse (Fig 5B and D). These observations indicate that APP/PS1 mice exhibit defective social recognition that can be corrected with intranasal oxytocin treatment (Fig. 5B to D).

**Fig. 5.**
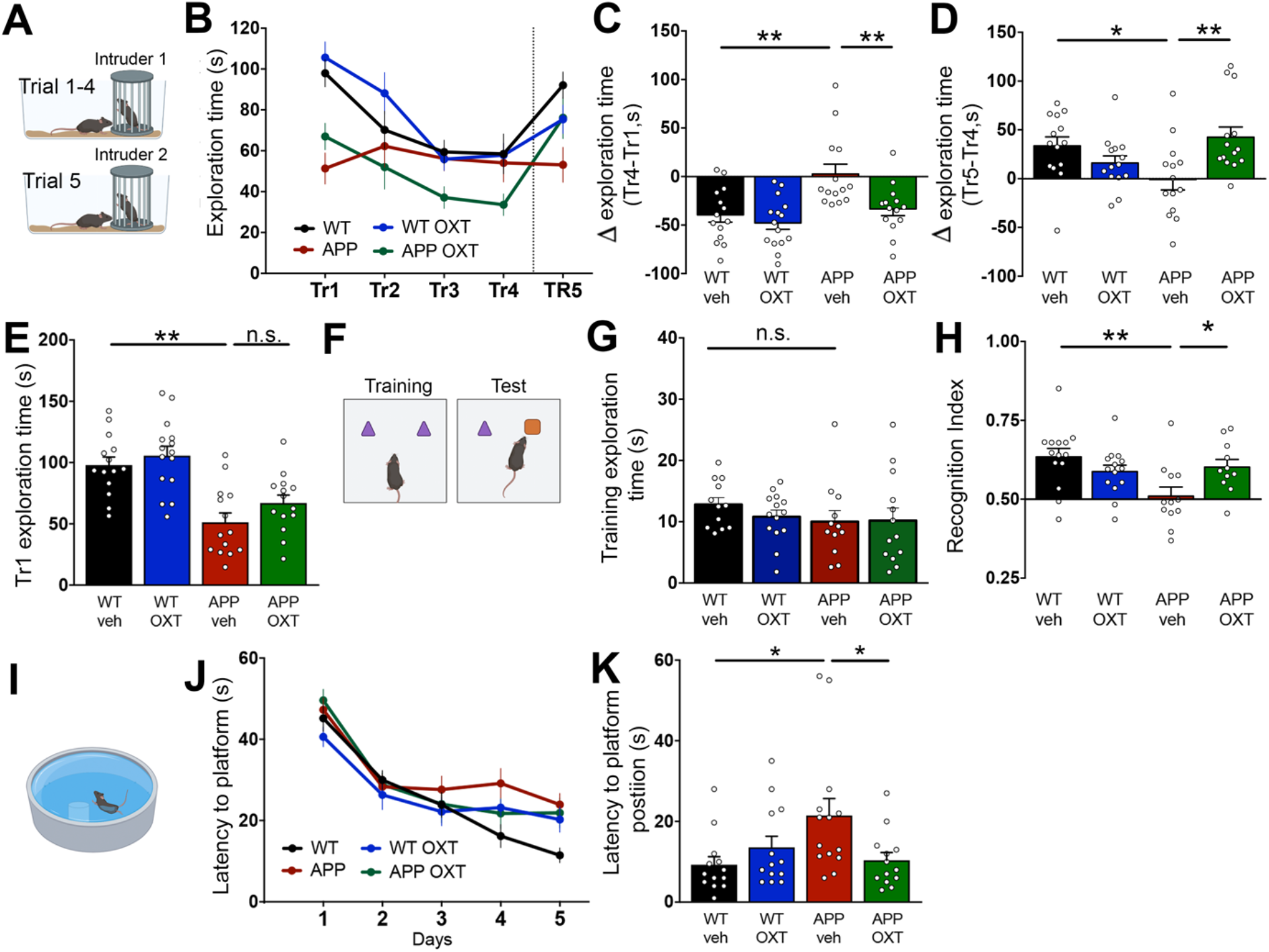
Intranasal oxytocin reverses memory deficits in aged male APP/PS1 mice. (**A**) Wild-type (WT) or APP/PS1 (APP) mice (10 months old) treated with intranasal vehicle (veh) or oxytocin (OXT) were tested in the 5-trial social test. The same stranger mouse was used in Trials 1 to 4. A novel stranger mouse was introduced in Trial 5. Time exploring the stranger mouse was quantified at 5-minute intervals along each trial. (**B**) Total time spent exploring the stranger mouse in each of the 5 trials; (**C**) Difference in exploration time between trials 4 and 1; (**D**) Difference in exploration time between trials 5 and 4; (**E**) Total exploration time in the first trial. Two-way ANOVA followed by Dunnett’s post-hoc test. (**F**) Mice were evaluated in the Novel Object Recognition task. (**G**) Total exploration time of the two objects in the 5 min training phase of the Novel Object Recognition task. (**H**) Short-term memory evaluated using the Novel Object Recognition task. Recognition index calculated as time exploring the new object/total exploration time. Values higher than 0.5 indicate novelty preference. (**I**) Mice were tested in the Morris Water Maze. (**J**) Latencies to reach the hidden platform in the training sessions. (**K**) Probe trial latencies to reach the area where the platform used to be. Two-way ANOVA followed by Dunnett’s post-hoc test. *p<0.05, **p<0.01.

Because oxytocin treatment had no effect on the decreased social exploration time exhibited by APP/PS1 mice during the first trial in this 5-trial test (Fig. 5B and E), we hypothesized that this could reflect a general reduction in exploratory behavior in APP/PS1 mice. To address this possibility, we quantified the total amount of time mice spent exploring the two objects presented in the 5-min training phase of a novel object recognition task (Fig. 5F), and found no differences in total exploration times comparing APP/PS1 and wild-type mice (Fig. 5G). This suggests that the reduced exploration time exhibited by APP/PS1 mice in the 5-trial social task is not caused by a generalized reduction in exploratory activity but rather reflects decreased sociability. Collectively, results from the 5-trial social recognition test indicated that oxytocin treatment corrected social memory deficits but did not rescue sociability in APP/PS1 mice. Our findings thus suggest that oxytocin could be protective against social memory loss in AD.

Oxytocin has been shown to regulate hippocampal memory consolidation (*28*) and to prevent Aβ-induced impairment in synaptic long-term potentiation (*29*). We thus hypothesized that oxytocin treatment could also rescue non-social memory impairments in APP/PS1 mice. Remarkably, intranasal oxytocin treatment restored memory of aged APP/PS1 mice in both the novel object recognition test (Fig. 5H) and in the Morris water maze (MWM) test (Fig. 5I and K). Control measurements showed that oxytocin had no effect on the MWM learning curve (Fig. 5J). Together, these results show that oxytocin not only rescues social memory deficits in the APP/PS1 AD mouse model, but it also protects against other types of non-social memory, including object recognition and spatial memory.

## Discussion

Despite significant efforts in pre-clinical research and clinical trials, no drugs to date have demonstrated clear clinical benefits to AD patients. Repurposing currently approved drugs may comprise a faster, safer and cost-effective pathway to the development of novel treatments for AD. Moreover, while the majority of therapeutic approaches tested in AD models use spatial and recognition memory tasks as readouts of efficacy (*30*), it is important to note that AD severely impacts social memory (*31*). The effects of potential therapies on social memory have been much less studied despite the fact that the inability of AD patients to recognize family members and friends is one of the biggest emotional burdens for their relatives. Here, we evaluated the therapeutic potential of the pro-social, anti-inflammatory hormone, oxytocin, using social and non-social memory tasks in APP/PS1 mice.

Based on our initial observation that *in vitro* and *in vivo* AD models presented reduced hypothalamic expression of oxytocin, we developed a protocol for chronic intranasal treatment with oxytocin. Intranasal administration of oxytocin is a translational, non-invasive route to deliver oxytocin to the brain; however, results in the literature are quite variable. This could be due, among other reasons, to use of different mouse strains, ages, sex and experimental protocols employed (*32-34*). We found that our experimental protocol effectively increased oxytocin peptide levels in the hippocampus and decreased fear response, an effect that has been described for central oxytocin (*18*). Oxytocin is approved by the US Food and Drug Administration for intramuscular or intravenous use (NDA #018248), and intranasal administration of oxytocin is currently being tested in clinical trials as treatment for obesity (NCT03043053), post-traumatic stress disorder (NCT03238924, NCT01466127), age-related inflammation (NCT02069431) and frontotemporal dementia (NCT01386333, NCT01937013, NCT01002300, NCT03260920). However, to date there are no studies registered in the ClinicalTrials.gov database to evaluate the actions of oxytocin in AD.

Our results show that treatment with non-invasive, intranasal oxytocin attenuates microglial activation, modulates plaque deposition by favoring formation of dense core plaques, and corrects memory deficits in APP/PS1 mice. Mechanistically, the promotion of dense core plaque formation by oxytocin may, at least in part, underlie its beneficial actions, as sequestration of soluble Aβ species into compact, dense core plaques has recently been suggested as a key mechanism by which microglia protect the brain from AD-related degeneration (*25*) and unpublished data from our group shows that neutralizing soluble Aβ oligomers rescues memory deficits in APP/PS1 mice. In line with a recent paper showing protective effects of oxytocin in early AD (*14*), our study shows for the first time that oxytocin can modulate inflammation and rescue social and non-social cognitive impairment in aged AD mice. These results pave the way for more detailed investigation of the cellular mechanisms and functional consequences of oxytocin signaling in AD. Importantly, considering that oxytocin is a commercially available endogenous hormone, safe and approved for clinical use by the US Food and Drug Administration and other regulatory agencies, our findings suggest that repurposing oxytocin could comprise a novel therapeutic strategy for AD.

## Materials and Methods

### Preparation and characterization of AβOs

AβOs were prepared from synthetic Aβ_1–42_ (California Peptide), as previously described (*35,36*), and were characterized by size exclusion HPLC and, occasionally, by Western blots using NU4, an oligomer-sensitive monoclonal antibody (*37*). AβO preparations comprise a mixture of Aβ dimers, trimers, tetramers, and higher molecular weight oligomers (*38*). Protein concentration was determined using the BCA assay (Thermo-Pierce).

### Animals

Three-month-old male C57BL/6 and Swiss wild-type mice were used for hypothalamic slice cultures and cannula implantation, respectively. Male 12-13 month-old APP/PS1 (or wild-type littermates) mice were used to determine oxytocin mRNA levels in the hypothalamus, and 9-10 month-old APP/PS1 (or wild-type littermates) mice were used for intranasal oxytocin treatment. Animals were housed in groups of five per cage with free access to food and water, under a 12 h light/dark cycle with controlled room temperature and humidity. All procedures followed the Principles of Laboratory Animal Care from the National Institutes of Health and were approved by the Institutional Animal Care and Use Committee of the Federal University of Rio de Janeiro (protocol #IBqM 050/20) and the Institutional animal care and use committee (IACUC) for NYU Langone Health (protocol #IA16-01096).

### Hypothalamic slice cultures

C57Bl/6 mice (3 month-old) were euthanized by cervical dislocation. Their brains were removed and hypothalami were dissected and sectioned in 400 μM slices using a McIlwain Tissue Chopper. Slices were exposed to vehicle or 500 nM AβOs for 3h after 1h of incubation in aCSF. After treatment, samples were collected for qPCR analysis of oxytocin mRNA levels.

### Implantation of intracerebroventricular cannula

This protocol was adapted from Boschi et al. (1981) (*39*). Briefly, 3 month-old Swiss mice were anesthetized with ketamine(100mg/kg)/xylazin(10mg/kg and placed in a stereotaxic frame. The coordinates used to implant a cannula in the lateral ventricle were: -0,6 mm Antero-Posterior; - 1,2 mm Medial-Lateral e -2,6 mm Dorsal-Ventral. (Allen Brain Atlas). Acrylic cement and stainless steel screws were used to fix the cannula to the bone. After implantation, mice were allowed to recover for 5 days and then received 5 consecutive daily injections of 10 pmol (in 3 μL) AβOs or vehicle.

### RNA extraction and quantitative real-time PCR

RNA was extracted from homogenized samples with the SV total RNA isolation kit (Promega) or RNAeasy Minikit (Qiagen). mRNA purity was determined using the 260/280 nm absorbance ratio and one µg mRNA was used for cDNA synthesis using the High-Capacity cDNA Reverse Transcription Kit (Applied Biosystems) or QuantiTect Rev. Transcription Kit (Qiagen). PCR was performed on an Applied Biosystems 7500 RT–PCR system or StepOnePlus™ Real-Time PCR System using the Power SYBR kit (Applied Biosystems). Fold changes in gene expression (2^-^ ΔΔ^Ct^) were calculated using cycle threshold (Ct) values (*40*). The following primers were used to detect oxytocin mRNA: Forward: GAGGAGAACTACCTGCCTTCG; Reverse: TCCCAGAAAGTGGGCTCAG; actin-β was used as a normalizer, using the following primers:Forward: 5’TGTGACGTTGACATCCGTAAA3’; Reverse: 5’GTACTTGCGCTCAGGAGGAG3’.

### Intranasal administration of oxytocin

Mice were manually and gently restrained and received 10 μL of a solution containing oxytocin or vehicle. This procedure was repeated 3 times a week for 6 weeks. The optimal dose of oxytocin used was determined in a pilot experiment in which mice received 3 different doses of oxytocin (0.8 μg, 8 ng and 8 pg) 3 times a week for 4 weeks prior to an intracerebroventricular injection of AβOs, followed by evaluation of cognitive impairment using the Novel Object Location test. The dose selected for subsequent studies (8 ng) was found to protect mice from AβO-induced memory loss and did not *per se* induce any cognitive impairment.

### Oxytocin ELISA

Oxytocin was quantified in the hippocampi of animals treated with vehicle or oxytocin using an ELISA kit (Enzo) and following manufacturer instructions. Briefly, hippocampi were dissected from the brains after cervical dislocation, and were homogenized in PBS containing protease and phosphatase inhibitor cocktail (Thermo Scientific Pierce). After sample centrifugation, supernatants were used for oxytocin determination.

### Contextual fear conditioning

The test was performed as previously described (*41*), with the following adaptations: In the training phase, mice were presented to the conditioning cage (40 × 25 × 30 cm), which they were allowed to freely explore for 2 minutes followed by application of two sequential foot shocks (0.8 mA) spaced by 30 sec. Mice were kept for another minute in the cage and removed. In the next day, mice were presented to the same cage for 5 minutes, without shocks. Freezing behavior was recorded automatically using the Freezing software (Panlab; Cornella, Spain) and was used as a fear memory index.

### Immunohistochemistry

Brains were fixed by perfusing anesthetized animals with saline, followed by 4% paraformaldehyde in phosphate-buffered saline (PBS). Brains were removed, cryoprotected using increasing concentrations of sucrose, frozen in dry ice and stored at -80 °C. Brains were subsequently sectioned (40 µm) in a cryostat (Leica Microsystems) and stored in PBS. After washing with PBS, coronal sections were blocked for 2 hours with 0.3% Triton X-100 and 5% BSA in PBS at room temperature, and incubated overnight a 4 °C with anti-oxytocin (1:500, Millipore), anti-Iba1 (1:200; Wako or Abcam) or 6e10 (1:1,000; Biolegend) primary antibodies diluted in blocking buffer. On the following day, sections were washed and incubated for 2 h at room temperature with Alexa Fluor 555-, 594- or 488-conjugated secondary antibodies (1:1,000; Abcam) and mounted with Prolong with DAPI (Invitrogen). Images were acquired using a Zeiss Axio Observer Z1 microscope, area and intensity levels were quantified on FIJI.

### Microglia isolation and culture

Microglial cells were isolated from P18 rat pups as previously described (*27*). Briefly, after tissue dissociation using DNAseI (Worthington) and myelin and debris removal, microglial cells were isolated by immunopanning with the OX-42 anti-Cd11b antibody (BIO-RAD). Cells were plated in 12-well chamber slides (Ibidi) and cultured during 5 DIV at 37°C and 10% CO_2_. A 50% media change was performed every 2 days.

### Immunocytochemistry

Microglial cultures were treated with AβOs (500nM) and/or oxytocin (10nM) for 3h. Cells were fixed with 4% paraformaldehyde in DPBS (Gibco) for 10 minutes. Cells were permeabilized with Triton X-100 diluted 0.1% in DPBS for 10 minutes and blocked using Normal Donkey Serum (NDS) 10% in DPBS for 1h at room temperature. Iba1 primary antibody (Abcam) was diluted 1:500 in blocking buffer and incubated at 4°C overnight. Secondary anti-goat antibody (Abcam) was prepared 1:1,000 in 1%NDS and incubated for 1h at room temperature. Cells were washed 3 times with DPBS between steps. Cells were mounted in Prolong with DAPI (Invitrogen) and imaged in a Zeiss LSM 800 confocal microscope. Iba1 intensity levels were quantified on FIJI and process number were quantified manually.

### Five-trial social test

The test was performed as previously described (*27,42*). Briefly, mice in experimentation were allowed to socially interact with a stranger mouse for 5 minutes during 4 sessions. In the 5^th^ and last session, the stranger mouse was replaced by a new one. The interaction times of the experimental mice with the introduced mouse in trials 1 to 5 were quantified.

### Novel object recognition task

This was performed as previously described (*38*). Briefly, an open field arena measuring 30 × 30 × 45 cm (W x L x H) was used. Before training, each animal was allowed to freely explore the arena for 5 min. During the training phase, the mice were placed in the arena in the presence of two identical objects which they could explore for 5 min. For the test, one of the objects was replaced by a novel one. The experiment was recorded and the recognition index (RI) was calculated as RI = time exploring the novel object / (time exploring the novel object + time exploring the familiar object).

### Morris Water Maze

This was adapted from Vorhees and Williams (2006) (*43*). Briefly, animals were placed in a pool with a hidden platform. During 5 days, mice were trained in 4 trials/day, beginning a different starting points. After each 60 second-long trials or once they reached the platform, the animals were allowed to remain for 40 seconds in the platform. Latencies to find the platform were quantified. Distance travelled and velocity were also measured. On the 6th day, the platform was removed for the probe trial, and the latency to reach the region where the platform used to be was quantified.

### Statistical Analysis

Statistical analyses were performed using the GraphPad Prism. Sample sizes and tests used are detailed in the figure legends.

## Acknowledgments

We thank Dr. Eric Klann and Dr. Martin Sadowski for donation of mice and NYU Langone Microscopy Core for experimental and technical support. We thank Dr. Luis Eduardo Santos and Jessica Minder for critical discussions.

## Author contributions

Conceptualization: MCS, RCF, MVC, STF. Methodology: MCS, JTSF, YPRF, BML. Funding acquisition: RCF, MVC, STF. Project administration: MCS, RCF, MVC, STF. Supervision: RCF, MVC, STF. Writing – original draft: MCS. Writing – review & editing: RCF, MVC, STF.

## Data and materials availability

All data are available in the main text or the supplementary materials.

